# Two new species of *Crocus* series *Scardici* from the Western Balkans

**DOI:** 10.1101/2022.10.27.514018

**Authors:** Donald Shuka, Helmut Kerndorff, Nomar Waminal, Vladimir Ranđelovič, Doerte Harpke

## Abstract

*Crocus* series *Scardici* has only two members, *C. scardicus* and *C. pelistericus* distributed at higher elevations in the Balkan Peninsula. In the course of our research, we discovered two new species growing on alpine meadows and pastures in the serpentine massifs in Albania and limestones in North Macedonia, respectively. The morphology and phylogenetic analyses (two nuclear single-copy markers, nrITS, two chloroplast markers) place the new species within series *Scardici*. Both new species are closely related to *C. scardicus*, but morphologically and ecologically clearly differentiated. We also measured genome sizes for all series *Scardici* species and report here the largest genomes found up to now in crocuses. The impact of different habitat types and isolation on separate mountain ranges in this alpine group of plants is discussed, too.

## Introduction

Series *Scardici* Mathew has only two members, *Crocus scardicus* Košanin and *Crocus pelistericus* Pulević, endemic to the western Balkans. Due to their possession of prophylls, they were sorted into section *Crocus* by Mathew (1982). Both species can easily be distinguished from all other members of the genus by their lack of a white stripe in the middle of the upper side of the leaves (Mathew 1982). According to Ž. Sekovski and G. Dimeska (Ranđelovič et al. 2007) both species share the same chromosome number 2n = 34.

Morphologically, *C. scardicus* has a unique bicolored flower with a lower purple and an upper yellow to orange part while flowers of *C. pelistericus* are characterized by a deep violet. Both are alpine species with a limited distribution range within the Scardo-Pindian (Scardian Mts.) subregions of the Middle European floristic region (Kuzmanović et al. 2016). *Crocus scardicus* occurs in the area of the Sharri, Kallabak, and Korabi mountain ranges at the borders between northeastern Albania, southern Kosovo and western parts of North Macedonia (Ranđelovič et al. 1990; Shuka et al. 2009; Hashani et al. 2014) with its southern distribution limit on Mt. Jablanica (Ranđelović et al. 2012). *Crocus pelistericus* thrives in the alpine areas of Pelister and NidŽe in North Macedonia and Kajmakčalan, Voras and Varnous mountains in northern Greece, growing in damp alpine turf.

Several of the mountains within the species’ distribution ranges are remote and largely unexplored. Moreover, the border regions of Albania have been considered as “forbidden terra” between 1945-1992. Therefore, it is not surprising that frequently new species have been found there during last 15 years (e.g., Shuka & Jahollari 2007; Shuka 2008, Shuka & Tan 2009; Ranđelović et al. 2012; Wagenitz et al. 2018; Raca et al. 2020).

*Crocus scardicus* and *C. pelistericus* prefer siliceous soils at elevations between 1700 m and 2450 m. According to Kilias et al. (2010) this metamorphic belt developed during the Paleocene to Eocene and stretches between the Korabi-W. Pelagonian zone up to the east of it in the Vardar/Axios ophiolitic belt. According to Čarni & Matevski (2010) the habitats of *C. pelistericus* are oligotrophic alpine mire with tussocks of *Sphagnum* spp. and sedges (*Carex* spp.) or transition mires and quaking bogs, characterized by acidic fen formations dominated by *Carex macedonica*. In contrast, *C. scardicus* prefers semi-arid or dry alpine pastures and meadows vegetated by associations of *Seslerion comosae* Simon, *Poion violaceae* Ht. and *Ranunculion crenati* Lakusić often dominated by *Nardus* grasslands and violets of sect. *Melanium* on siliceous substrates. Some subpopulations of the species can be found also in the areas where limestone overlaps with siliceous soils (Randjelović et al. 1990; Hashani et al. 2014; S∤omka et al. 2014). Reports on the presence of *C. pelistericus* in Albania (Barina et al. 2015) and Jablanica Mt. in North Macedonia (Rukšāns & Zubov 2022) refer to the new species described here.

On the Jablanica mountain range one new member of Series *Scardici* was found by Dörte Harpke together with Novica and Vladimir Ranđelovič in June 2013 during an excursion and recognised as a new species according to morphological peculiarities and differences in molecular data compared to both of its relatives. It is growing adjacent to *C. scardicus* there but has pale violet flowers. Several attempts to find this new crocus in flower again to carry out additional morphological investigations failed in the following years until May 2019.

The other new species of Series *Scardici* was identified in the serpentine massifs between the municipalities of Gramsh, Maliq and Pogradec in southeast of Albania. During a floristic investigation of the Valamara-Guri i Topit mountain range in June 2019, Donald Shuka found an alpine meadow colored yellow by a *Crocus* species in bloom. Different populations were investigated by D.S. during 3 years in the area bordered by Lenia Mt. in the south and the Lakes of Lukova in the north of Valamara Mt. range. Morphologically it is similar to *C. scardicus* but having a white or yellowish lower part of the flower or completely white flowers. Occasionally, also in populations of *C. scardicus* individuals with a lower white instead of violet part of the flower can be found. However, there are additional morphological characters that differentiate it from *C. scardicus*.

Within this study, we attempt to (*i*) characterize the morphology of both new species, and (*ii*) infer their affiliation with series *Scardici*. Since molecular marker sometimes failed to fully resolve phylogenetic relationships within the genus, we investigated here five molecular markers comprising nuclear as well as plastid regions.

## Material and Methods

### Molecular analyses

We included nine populations each with 1-3 individuals representing four series *Scardici* species and 28 outgroup taxa for internal transcribed spacer (ITS) region of the nuclear ribosomal DNA (nr DNA), a nuclear single-copy gene marker of the *ORTHO04×5M009129* orthologous gene family (orcp; Nemati et al. 2019) as well as a nuclear single-copy marker of *ORTHO04×5M007634* orthologous gene family (embo; supplementary table). As plastid marker, a region comprising the *trn*K intron, *rp*S16 and *trn*Q and their intergenic spacers as well as part of the *ycf*1 gene were investigated (Kerndorff & Harpke 2022).

Extraction of genomic DNA and amplification were conducted according to Nemati et al. (2019). PCR was performed with 1 U Phusion High-Fidelity DNA Polymerase (Thermo Scientific) in the supplied Phusion GC Buffer, 200 µM of each dNTP, 0.5 µM of each primer, and about 20 ng of total DNA in 50 µl reaction volume in a GeneAmp PCR System 9700 (Perkin-Elmer). Amplification was performed with 3 min initial denaturation at 95 °C and 35 cycles of 30 s at 95 °C, 25–60 sec at the marker-specific annealing temperature (ITS: Blattner 1999; orcp: Nemati et al. 2019; chloroplast marker: Kerndorff and Harpke 2022; embo: table1) and 30 s at 70 °C, followed by a final extension for 8 min at 70 °C.

**Table.**
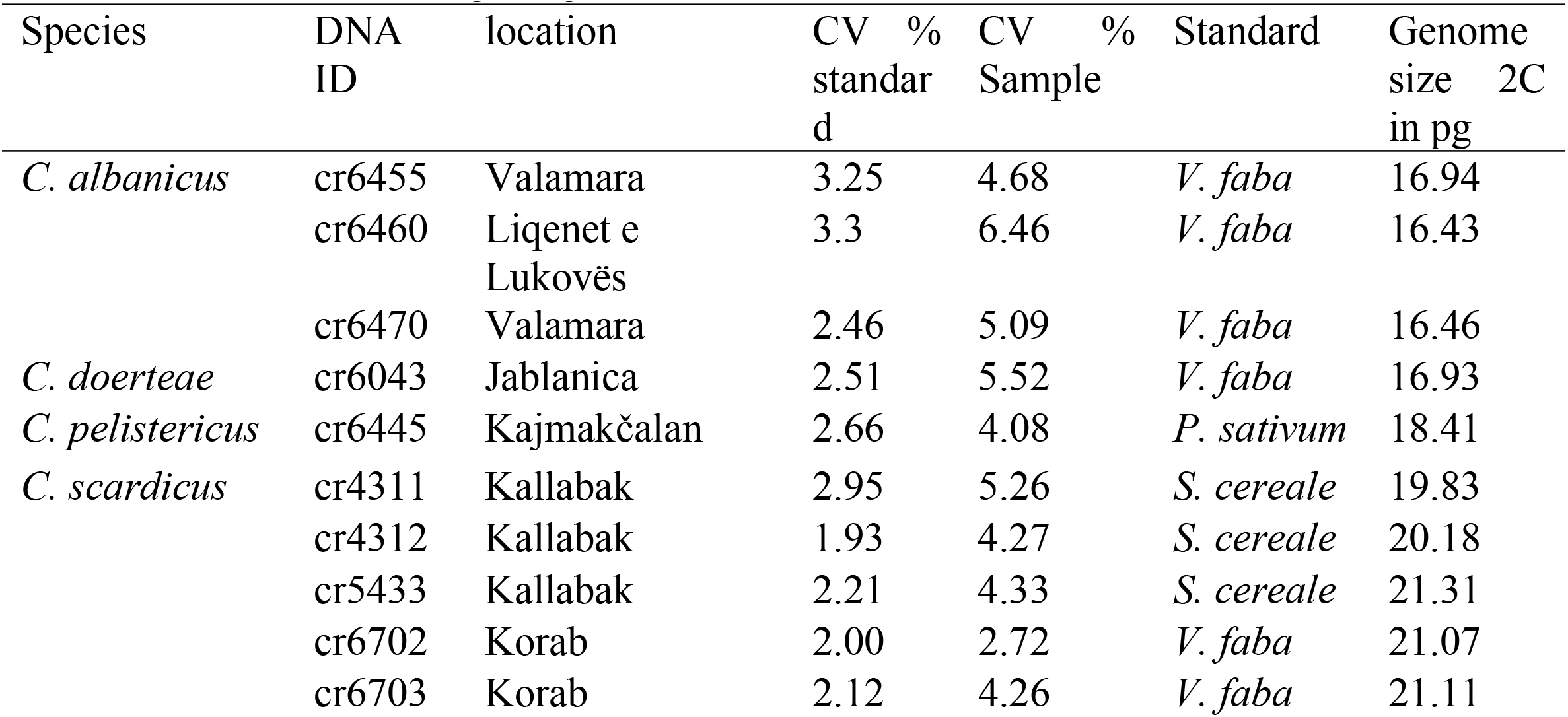
Genome size measuring using *Pisum sativum* as standard.

All newly obtained sequences were submitted to the ENA nucleotide database. Sequence accession numbers for the studied individuals are accessible via the study ID PRJEB44177. Sequences were manually aligned.

Mafft (Katho et al. 2002) was used to generate initial alignments for the loci, which were corrected manually where necessary. The sequences of the nrITS region and the nuclear single copy loci were concatenated resulting in one dataset. A second alignment resulted from the concatenation of the *mat*K-*trn*Q region as well as of *ycf*1. Herbarium vouchers are deposited at the herbarium of the IPK Gatersleben (*C. doerteae* GAT0034720, GAT0072606; *C. pelistericus* GAT0046107, *C. albanicus* GAT0083264-GAT0083267, GAT00832271-GAT0083274, *C. scardicus* GAT0034718, GAT0034719, GAT0083175).

### Phylogenetic analysis

Each data set was first analyzed separately to check for incongruences. The data sets for each marker as well as the combined data set of nuclear markers were subjected to Bayesian phylogenetic inference (BI) with MrBayes 3.2.6 (Ronquist et al. 2012). For BI two times four chains were run for 1.5 million generations under the GTR+Gamma+I model of sequence evolution, sampling a tree every 1000 generations. Converging log-likelihoods, potential scale reduction factors for each parameter and inspection of tabulated model parameters in MrBayes suggested that stationarity had been reached. The first 25% of trees of each run were discarded as burn-in. Two independent runs of BI analysis were performed to confirm that separate analyses converged on the same result. In each of the two analyses the same topology and similar posterior probabilities (pp) of nodal support resulted.

### Morphological analysis

Twenty to 30 individuals per population were subjected to morphological analysis. Characters of the flowers including stylus and anthers were measured on herbarized specimens. Counting of leaves, characterization of colors was done on wild living material.

Leaves of the crocuses investigated were initially stored in 70% Ethanol and afterwards cut with a microtome to thin slices of 75 μm and stained with “Astrablau” and Chrysoidine. Cut samples were transferred into water and washed thoroughly to receive their natural form.

### Calcium-oxalate crystals in the corm tunics

For calcium-oxalate crystals, we used the same treatment as Wolter (1990) for the preparation of the corm tunics. They were visualized with a Dino-Lite Edge Digital Microscope using polarized light and photo-stacking technique in cases where necessary. In these cases, no scale bars can be shown, but enlargements are in the same order of magnitude as the pictures with scale bars. The Ca-oxalate monohydrate crystals have a high birefringence index of nγ-nα = 0.16, which is near the one of calcite (0.172), means they can create strong interference colours, dependent on species. The form, size, localisation, and frequency of the crystals in the tunics can be different and are usable as an additional distinguishing parameter for crocuses.

### Genome sizes

Genome sizes of silica dried leaf material of two to three individuals were measured in a CyFlow (Partec) flow cytometer using PI as stain (Meudt et al. 2015), with *Vicia faba* L., *Secale cereale* L. or *Pisum sativum* as standard (*V. faba* 2C = 26.9 pg, *S. cereale* 2C = 16.19 pg, *P. sativum* 2C = 9.09; DoleŽel et al. 2007).

### Karyological analysis

Roots were cut about 2 cm from the tips and pretreated with 2 mM 8-hydroxyquinoline for 5 hrs at room temperature and then with cold water overnight in a refrigerator. The roots were then fixed in Carnoy’s Solution (3:1 ethanol-acetic acid) for 24 hrs and stored in 70% ethanol until use. Slides were prepared according to Waminal et al. (2012) and Rodríguez-Domínguez et al. (2017). Briefly, the root meristems were isolated by cutting about 2 mm of the whitish tip and cutting out the root cap. Then the meristems were digested with a pectolytic enzyme solution containing 2% Cellulase RS (C8003, Duchefa) and 1% Pectolyase Y-23 (P8004, Duchefa) for 1 hr at 40°C. The digested cells were suspended by adding ∼200 µL distilled water and vortexing for about 1 min, followed by centrifugation at 13,000 rpm for 7 min. The cells were fixed by adding ∼200 µL chilled Carnoy’s Solution followed by vortexing to resuspend the cells and then another centrifugation at 13,000 rpm to collect the cells. The supernatant was decanted and the pelleted cells were resuspended in aceto-ethanol (9:1 v/v) before mounting onto slides.

Slides were fixed in 2% formaldehyde solution (47608, Sigma-Aldrich) for 3 min and dehydrated in ethanol series (70%, 90%, 99%). Chromosomes were stained with 1 µg/µL DAPI in 2× SSC. Images were captured using a 100× objective of an Olympus BX61 fluorescence microscope (Olympus, Japan).

### Results and Discussion

Series *Scardici* forms a strongly supported monophyletic group (pp 1; Fig. 1) as sister to section *Crocus* taxa, which are either distributed on the Balkan Peninsula or westwards (e.g., C. *ligusticus, C. goulimyi, C. cambessedesii, C. versicolor, C. minimus, C. suaveolens, C. corsicus, C. serotinus* and relatives) and in the nuclear data set it also groups with series *Verni* representatives, *C. banaticus* and *C. malyi*. While this grouping is consistent with earlier phylogenetic investigations (five chloroplast markers: Petersen et al. 2008; *trn*L-F and nrITS: Harpke et al. 2013), we could here achieve better phylogenetic resolution and stronger support values.

**Figure 1.**
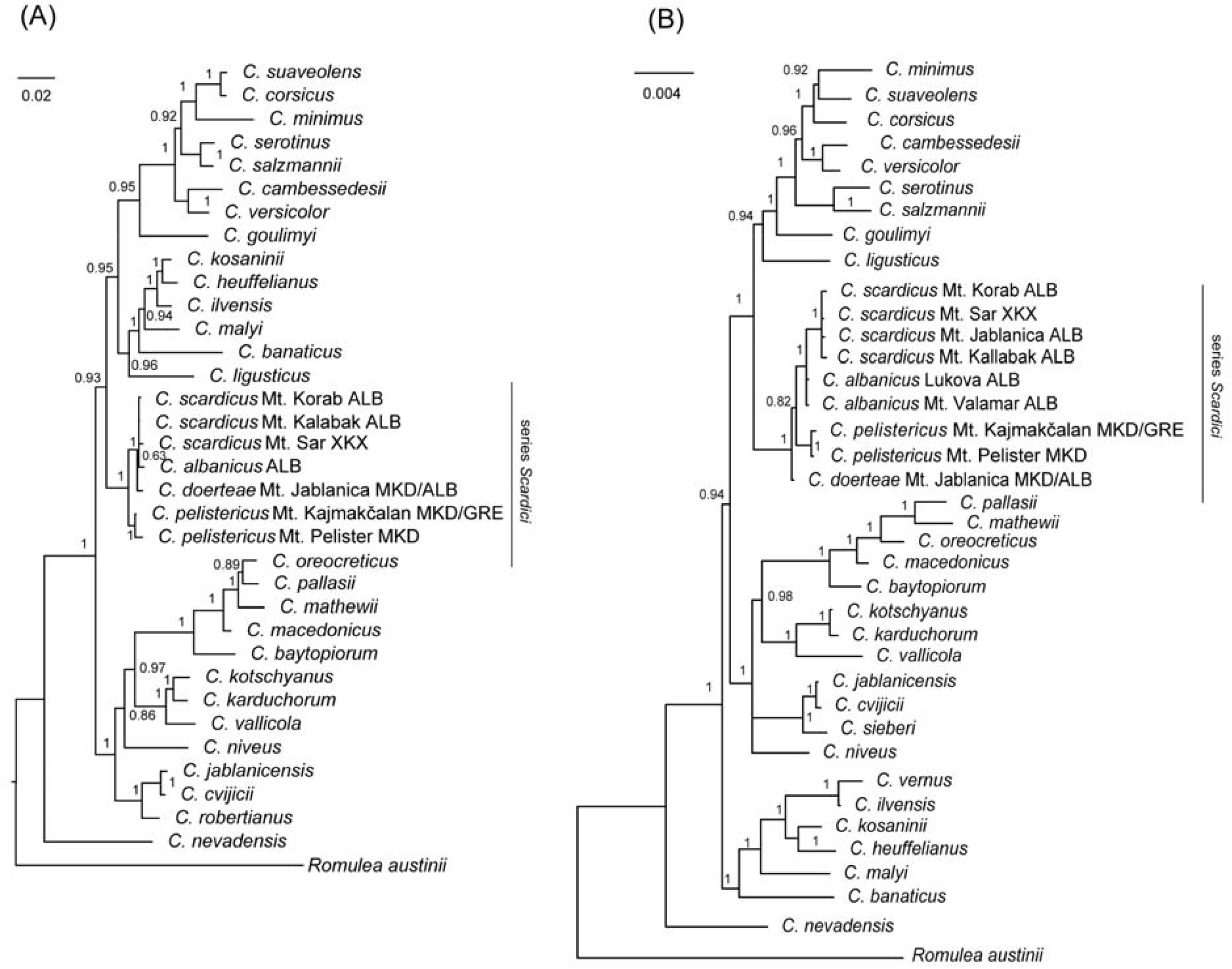
Phylogenetic trees derived from Bayesian phylogenetic inference based (A) on the concatenated alignments of embo, nrITS region and orcp (2489 bp) and (B) the combined chloroplast marker set of the concatenated *trn*K−*trn*Q plus *ycf1* regions (4514 bp). Posterior probabilities (pp) are given at the nodes of the tree.

Relationships within series *Scardici* differ a bit between the nuclear and chloroplast datasets. In the nuclear dataset *C. pelistericus* was found in a sister relationship (pp 1) to the other species whereas in the plastid data set one of the two new species, *C. doerteae*, is sister to the other species although with low statistical support (pp 0.88). In the nuclear data set, the latter species is grouped with the second new species, *C. albanicus*, and C. *scardicus* (pp 1). The difference between nuclear and plastid data set could be caused by incomplete lineage sorting as the result of (successive) rapid speciation events where an ancestral plastid genome was retained in *C. doerteae*.

The relationships between *C. albanicus*, and C. *scardicus* could not be resolved with the current set of markers albeit that there are differences (six substitutions in the *trn*K-*trn*Q region, embo: three substitutions, orcp: one substitution).

Initially, we measured genome sizes to identify potential differences in ploidy levels. The lowest genome size was observed in *C. albanicus* (2C = 16.43 – 16.94 pg; table) and *C. doerteae* (2C = 16.93 pg). *Crocus pelistericus* was measured with 2C = 18.41 (Tab. 2). The highest genome sizes were observed in *C. scardicus* (2C = 19.83 - 21.31 pg), where genome sizes variation is pronounced with up 1.5 pg difference within this species. However, our measurements of silica-dried material usually have higher CV values compared to measurements of fresh material, which might influence the accuracy of measurements.

Series *Scardici* crocuses possess the largest up to now measured genome of wild growing crocuses (see Raca et al. 2020; Kerndorff & Harpke 2022). Large genomes are frequently observed in alpine plants. It is hypothesized being connected with the presence of phosphate-rich soil at higher elevations supporting the high demand of phosphate for plants with large genome sizes (e.g., Albach & Greilhuber 2004; Guignard et al. 2017).

Morphologically, all series *Scardicii* members are characterized by the lack of the white leaf stripe. However, our cross sections of leaves show that the white stripe is usually not visible, because it is covered by several layers of chlorophyll-containing cells on top of the large area of mostly cells empty of chloroplasts, which normally are responsible for the appearance of a white stripe in the middle of *Crocus* leaves.

*Crocus doerteae* (Fig. 2) appears more similar to *C. pelistericus* by having violet flowers, but those are paler especially inside. Other distinguishing characters are the size of the flowers, length of perigone tube and filaments, width and number of leaves.

**Figure 2.**
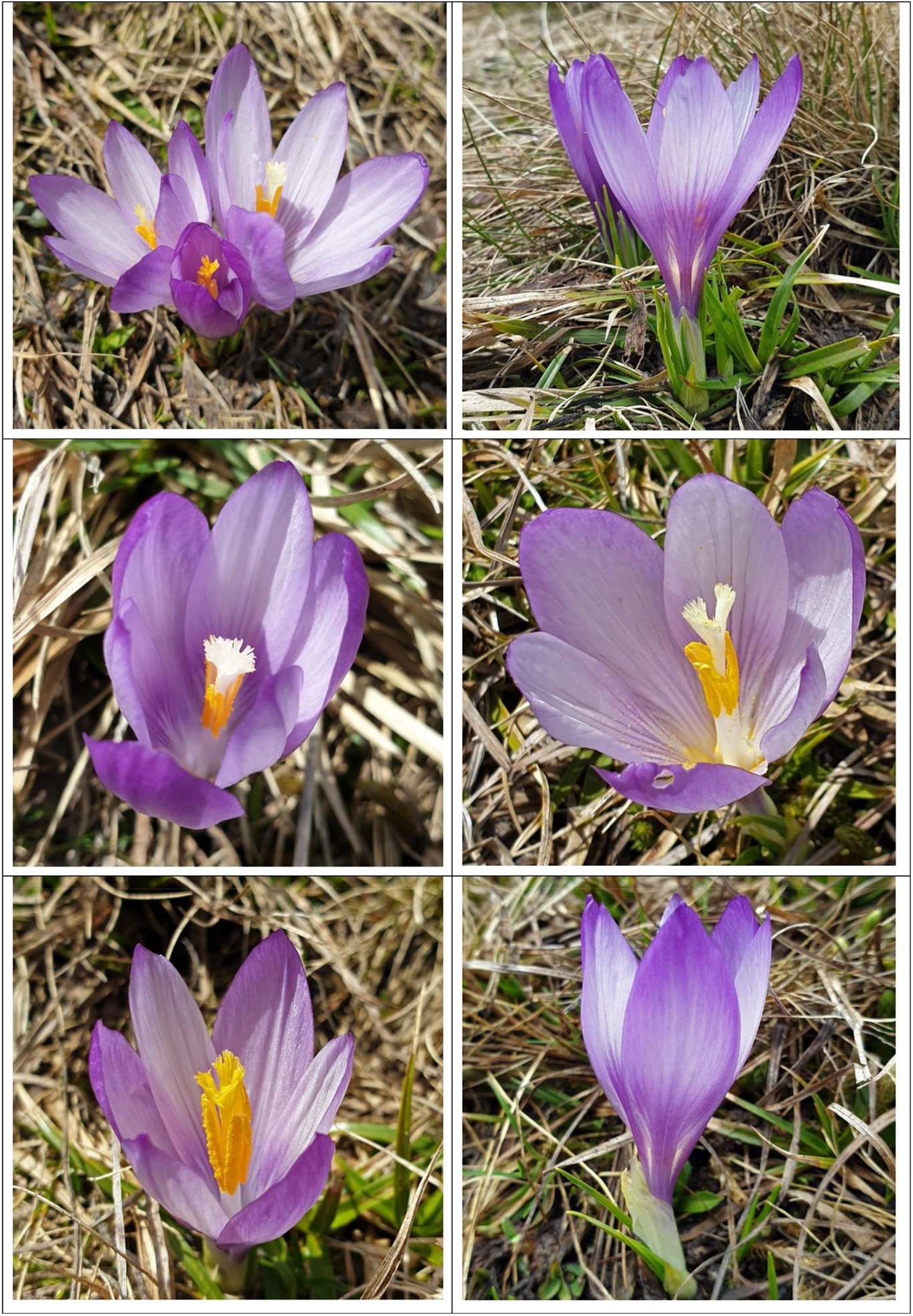
Habitus of *Crocus doerteae*.

Although *C. doerteae* is partly growing next to *C. scardicus*, we did not find evidence for current hybridization in the molecular or morphological data. *Crocus albanicus* is closely related to *C. scardicus*, with which it also shares a high morphological similarity. However, both species can be distinguished from each other by corm tunic fibers, throat color, shape and color of perigone segments, and leaf morphology.

*Crocus albanicus* (Fig. 3) differs from its closest relatives *C. scardicus* from the presence of white or yellowish throat of the flower (not purple), from its bigger flower and oblanceolate or elliptical perigone segments (not obovate), the wider leaves and their shape and habitat of growing, serpentine (not siliceous or limestone substrate). It is different also from *C. doerteae* regarding the flower color, the shape of leaves and perigone segments, as well as the color of tunics and shorter persistent neck (1.5–2 cm, not 2–4 cm such as to *C. doerteae*).

**Figure 3.**
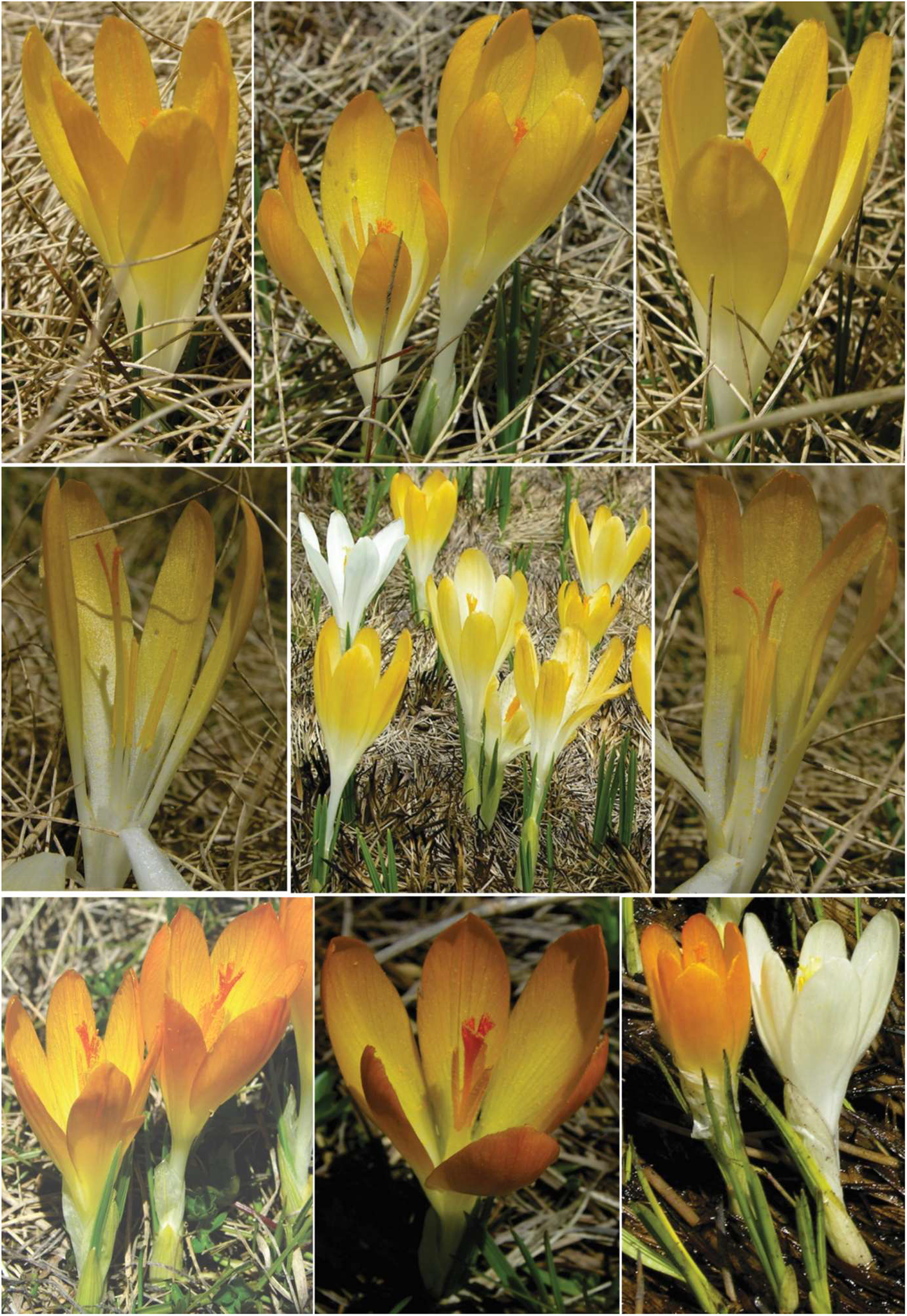
*C. albanicus*, color variations of flowers and the position of styles and stamen to each-others in flower.

The *Crocus* species of series *Scardici* all belong to the Scardo-Pindian floristic subregions and are endemics of Western Balkans, but differ in their growing habitat types and their distribution ranges. *Crocus albanicus* is a serpentine obligate species of the arid Mediterranean climate in alpine pastures and meadows of Valamara Mt. range, in contrast with *C. scardicus* which prefers siliceous or limestone substrates of the Alpine biogeographic region. They are growing allopatrically. The distribution area of *C. doerteae* is in the contact zone of calcareous rocks of the Cretaceous with ultramafic rocks of the Mirdita-Pindos ophiolites in between the distribution range of *C. albanicus* and *C. scardicus* (Fig. 4), partly overlapping with the latter species’ distribution. *Crocus scardicus* and *C. pelistericus* share almost the same siliceous substrate, but they occur in different habitat types with quite different ecological conditions. *Crocus pelistericus* occurs in wet or semi-wet alpine turf like *Crocus albanicus* and *C. doerteae*. However, these three species grow in different vegetation units with different pH, respectively ultra-alkaline fens, alkaline to neutral fens and acidic fens. All species of series *Scardici* are growing sympatrically with other crocuses (e.g., C. *albanicus* with *C. veluchensis* complex, *C. scardicus* with *C. veluchensis* complex, *C. doerteae* with *C. jablanicensis* and *C. veluchensis* complex, *C. pelistericus* with *C. veluchensis* and *C. sublimis* complexes).

**Figure 4.**
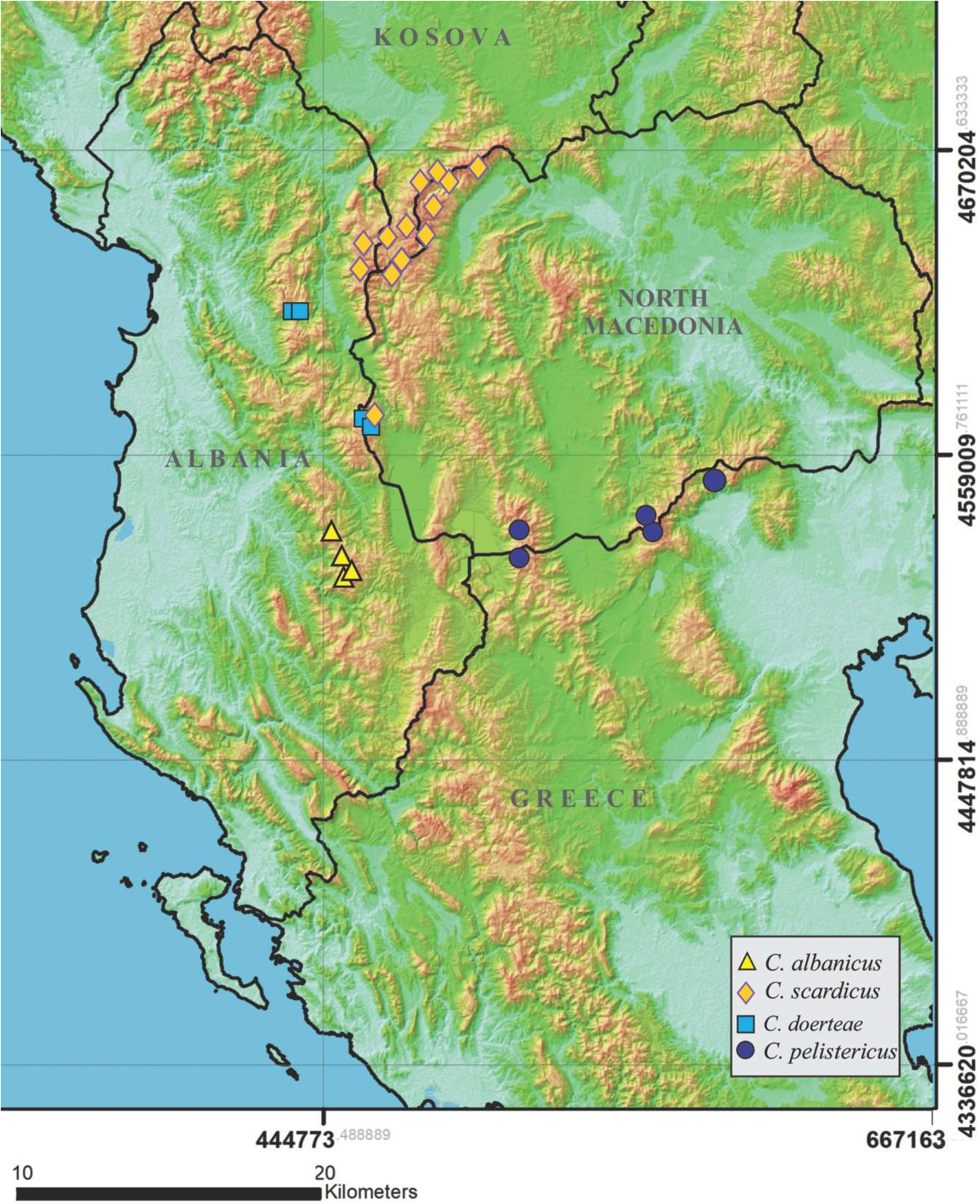
Distribution map of ser. *Scardici* in the western Balkans.

### Description of species

#### *Crocus doerteae* Kerndorff & V. Randjel. *sp. nov*. (Figs. 2, 5, 7)

Type:—NORTH MACEDONIA. Jablanica mountain ridge, on wet or semi wet meadows on limestone substrates, 18 June 2013, 1800-2100 m, *D. Harpke s*.*n*. (holotype: GAT0034720!, isotypes TIR!, GAT0034721).

**Figure 5.**
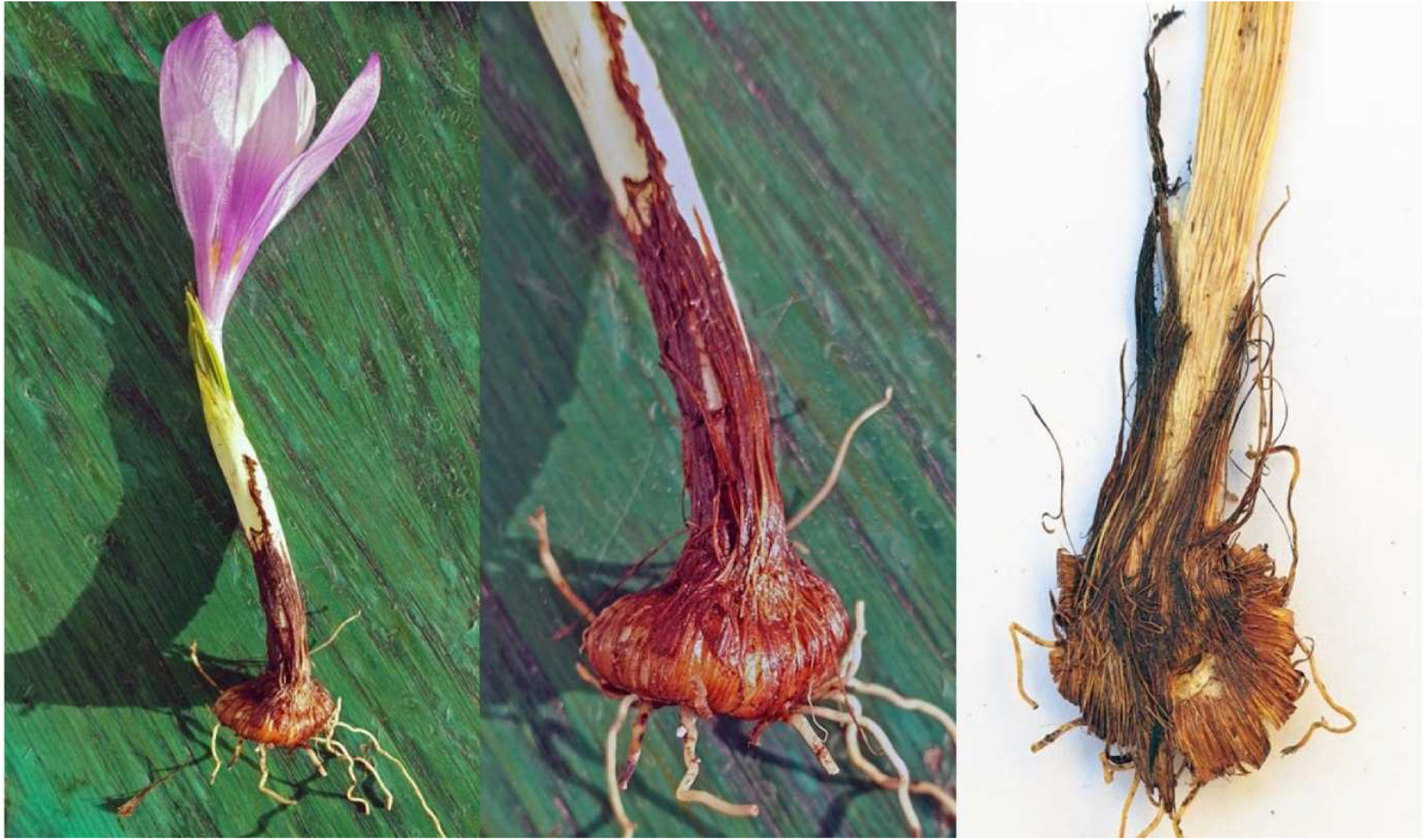
*Crocus doerteae* with flower, cataphyll, corm, tunic, and neck.

**Figure 6.**
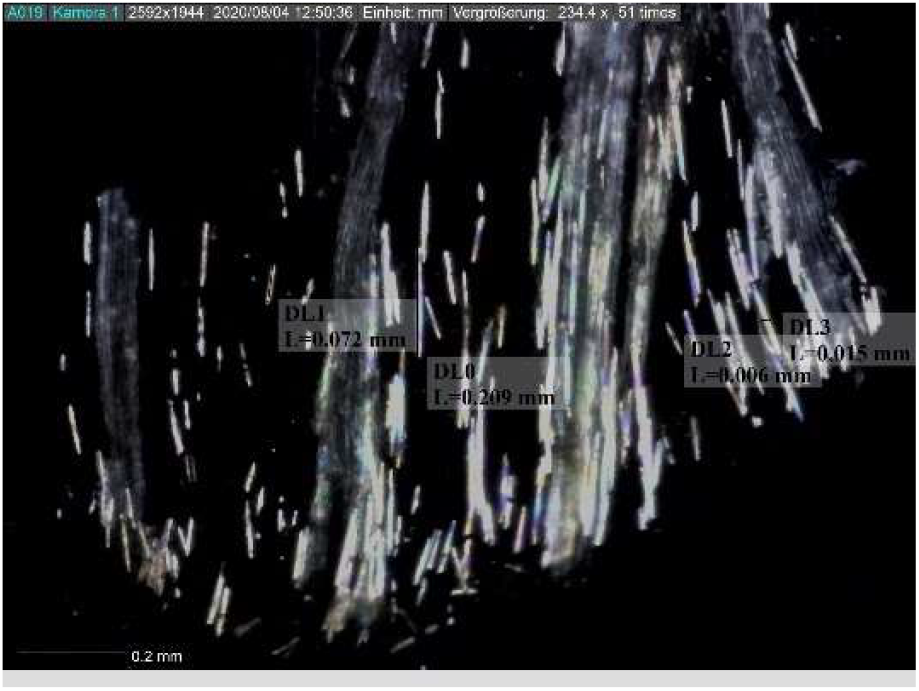
Oxalate crystals in the corm tunic of *Crocus doerteae*.

**Figure 7.**
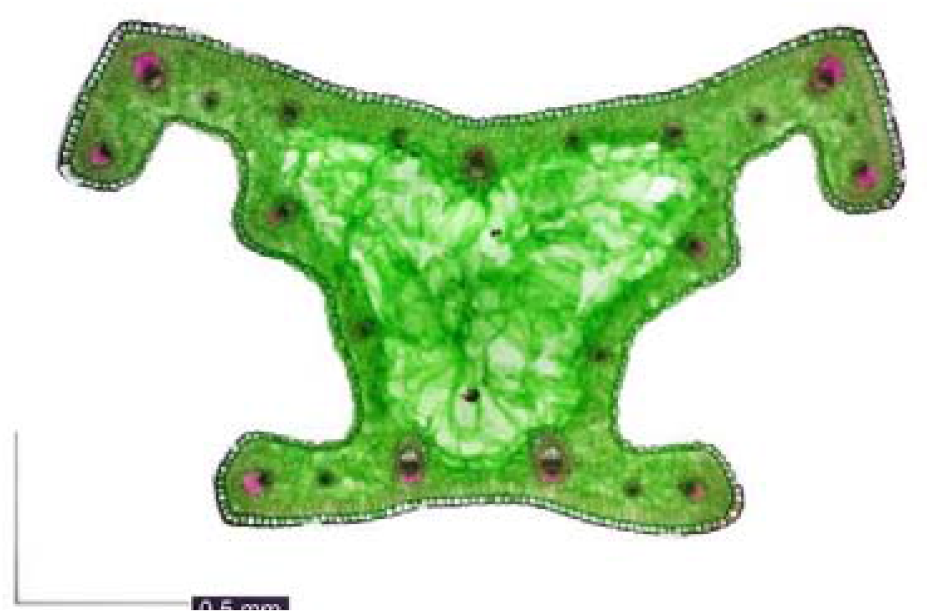
Cross leaf section *Crocus doerteae*.

The corm is very flattened globose with a diameter of 8–12 mm. Tunics fibrous and parallel approximately from the middle down, in the upper third fibers are slightly anastomosing merging into a long neck with parallel fibers which are partly connected with membranes. Due to permanent wet conditions in the soil the tunics are decayed comparatively quickly which results in only two, often incomplete tunic layers which are hardly to distinguish. The diameter of fibers is around 0.09 to 0.13 mm; Ca-oxalate crystals are unevenly distributed long needles of different size in several parts of the corm tunic, 0.006–0.015 mm wide and 0.072–0.3 mm long (Fig. 6). Cataphylls several, white to yellowish with green veins, at apex mostly brown and peaky. Leaves 3–**4.4**–7, light green to green, having one very peculiar shaped rib underneath of each leaf-arm. In contrast to most other crocuses, the white stripe is usually not visible, since several layers of chlorophyll-containing cells cover the large area of the mostly empty cells, which normally “creates” the “white stripe”; diameter of leaves 1.2–2 mm, glabrous. The leaf-basis (keel) is concave the arms are short and slightly open with an inclination of 10 degrees outwards (Fig. 7). Prophyll present. Bract and bracteole present subequal but not elongate at apex. White throat short, merging into the lilac-violet stripes of the segments, which become more intense from the middle down, glabrous. Perigone tube is violet, very short and mostly below 3 cm above ground. The inside of all segments is rather whitish-lilac, at apex more violet. The outside of the inner segments is similar. The outside of the outer segments is mostly deep violet, darker near perigone tube, sometimes with faint darker violet veins. Outer and inner segments are of different sizes. Outer segments are 29– **36**–41 mm long and 11–**14** (–16) mm wide (n = 23), inner segments are 27–**31** (–38 mm) long and 9–**11** (–12) mm wide (n = 23). Filaments white 9–**12.3** (–15) mm long glabrous (n = 22); anthers yellow to orange, 10.5–**12.3** (–14.5) mm long in the center of the apex with notch, anther lobes 1.5–**2.3** (–3.2) mm long (n = 22), connective rather broad, white, pollen deep yellow. Styles white and yellow in relation of about 50% (n = 25), with one level of branching, branching position in relation to the anthers mostly in the upper third of the anthers and above of them. The three short branches are expanded and fimbriated into very many thin and short threads at apex, length of style-branches including filiform ends 3–**5.3** (–10) mm (fig. 4). Length of styles according to stamen 19 (91%) longer, 2(9%) equal and 0(0%) shorter. Capsule 0.9–14 × 3–5 mm. Seeds 2–3.5 × 1.5–2 mm, dark reddish to blackish at maturity (Fig. 8: l, m). Chromosome number 2*n* = 34 (Fig. 8).

**Figure 8.**
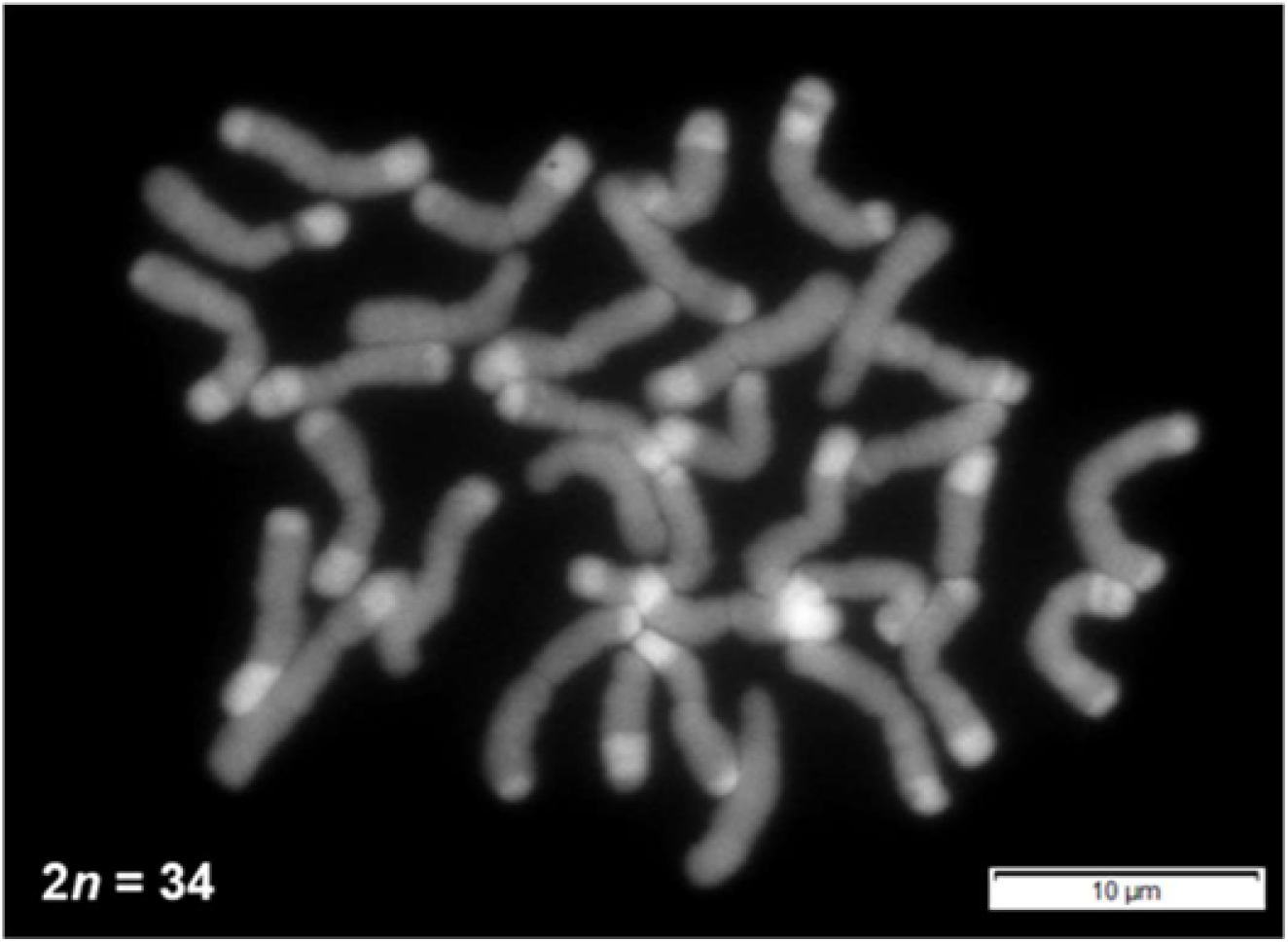
Chromosome count of *C. doerteae* roots showed 2*n* = 34 chromosomes.

### **Other specimen examined of *Crocus doerteae* (paratype)**:—

ALBANIA. Mati municipality, National Park Lurë-Mali i Dejës, at the foot of eastern side of Mali i Dejës, in meadow around Liqeni i Madh (Big Lake) on calcareous soil, 1900 m, 12 June 2019, *D. Shuka & L. Shuka s*.*n*. (herb. Shuka), TIR!).

—*loc. ibid*., 500 m below the Liqeni i Madh on serpentine pastures wet by the stream of water flowing from Liqeni i Madh, 1800 m, 10 June 2018, *L. Shuka s*.*n*. (herb. Shuka), identified as *C. pelistericus*.

### Habitat and Distribution

The distribution of *C. doerteae* is overlapping with *C. scardicus*, but grows also further south probably completely closing the distribution gaps between *C. pelistericus* and *C. scardicus*. It is like its relatives an alpine species growing between 1800 and 2100 m. Up to now the only known habitats are in N-Macedonia in the Jablanica mountain ridge in alpine turf and water-soaked source horizons over Paleozoic and Mesozoic carbonates (limestone formations). The second locality of *C. doerteae* was found in small wet alpine places on the eastern side of Deja Mt, northeast Albania, vegetated from mire communities that form peat. The locality occurs at the contact of calcareous rocks of Cretaceous with ultramafic rocks of the Mirdita-Pindos Ophiolites (Kilias et al. 2010). The species was found in alkaline fens of the *Caricion davallianae* community dominated by *Eriophorum latifolium, Carex davalliana, Scirpus holoschoenus* or *Dactylorhiza cordigera* and *Pinguicula balcanica*. The vegetation of springs and brooks on serpentine habitats, below the Liqeni i Madh include among others *Cardamine raphanifolia, Carex* spp., *Dactylorhiza cordigera, Geum coccineum, Parnassia palustris, Pinguicula balcanica, Ranunculus* sp., *Rumex alpinus, Veratrum album* or endemics such as *Schilla albanica* and *Ranunculus degenii*.

The flowering time is May and June, depending on the altitude and exposure.

### Etymology:—

The species name is dedicated to the botanist Doerte Harpke.

#### *Crocus albanicus* D. Shuka & Harpke, sp. nov

(Figs. 3, 9: a, b, c, d, e, f, 10: a, b, c, 11: a).

**Figure 9.**
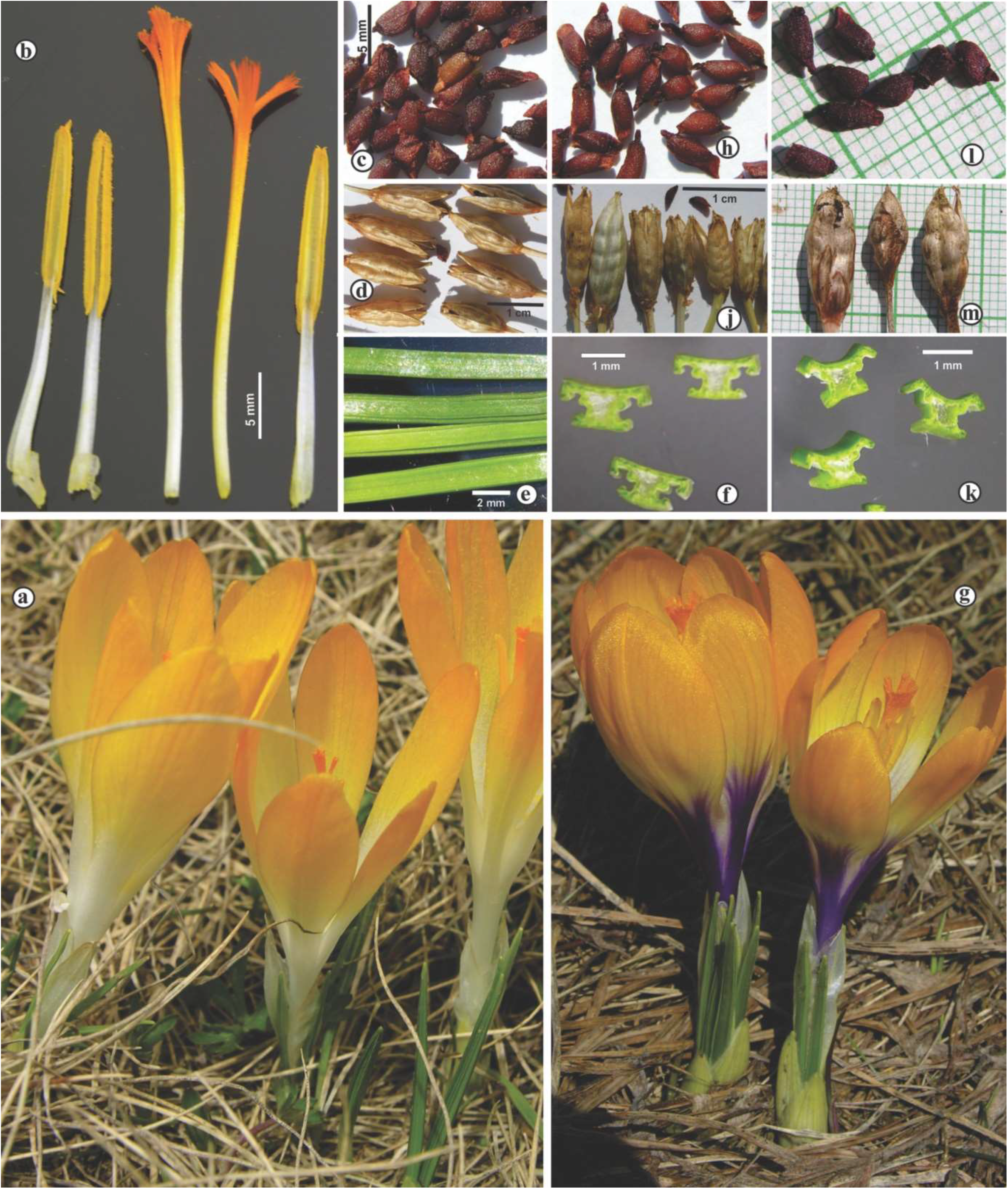
*Crocus albanicus*, a) habitus, b) style and anthers, c) seeds, d) capsules, e) leaves, f) cross section of leaves, g) *C. scardicus* habitus h) seeds, j) capsules, k) cross section of leaves, l) *C. doerteae* seeds and m) capsules.

**Figure 10.**
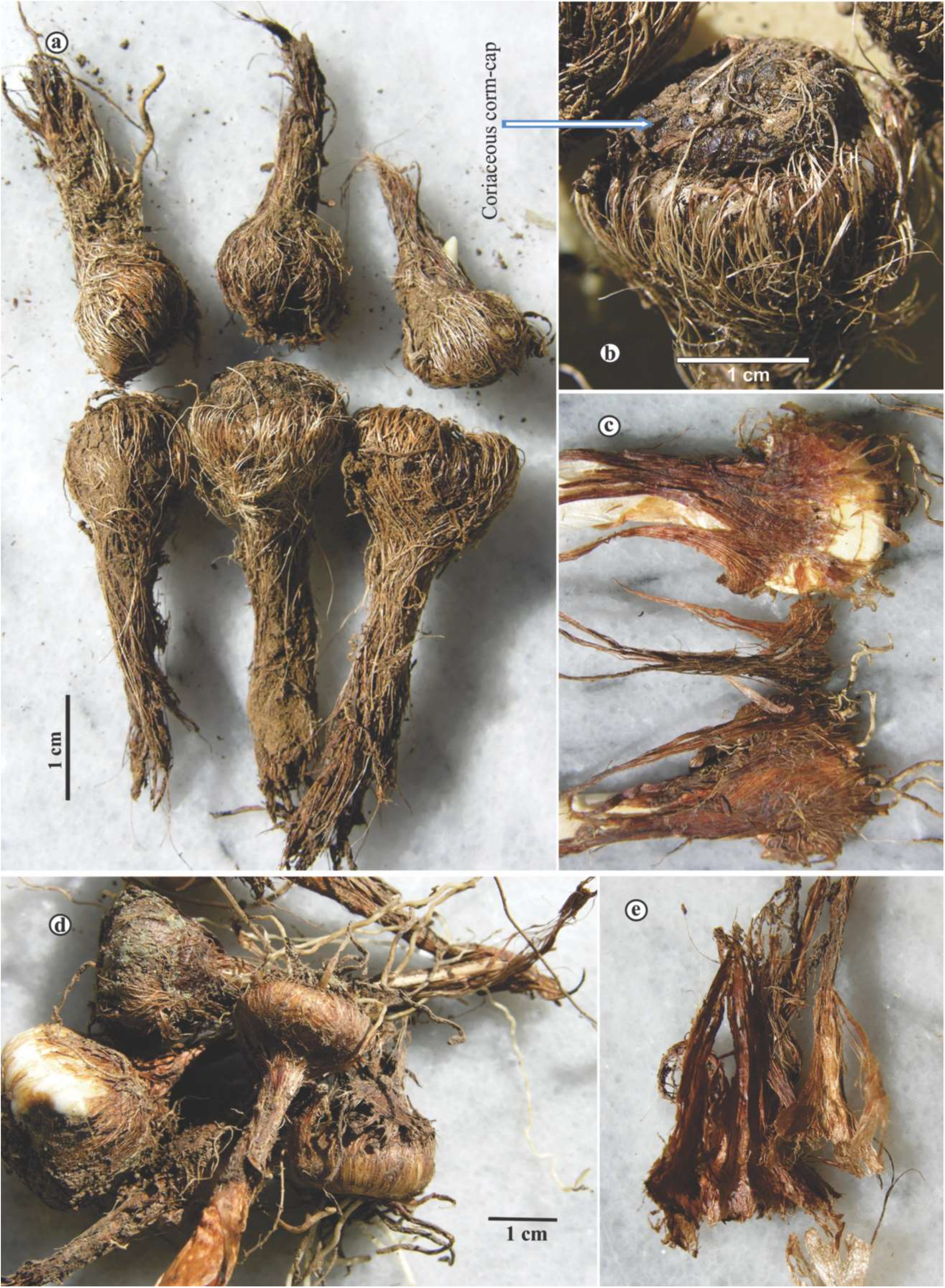
*C. albanicus* corms and tunics, a) corms with outer thick fibers, b) coriaceous corm-cap, c) inner membranous layers with thin fibers, d) *C. scardicus* corms and tunics, e) inner membranous layers with thin fibers.

**Figure 11.**
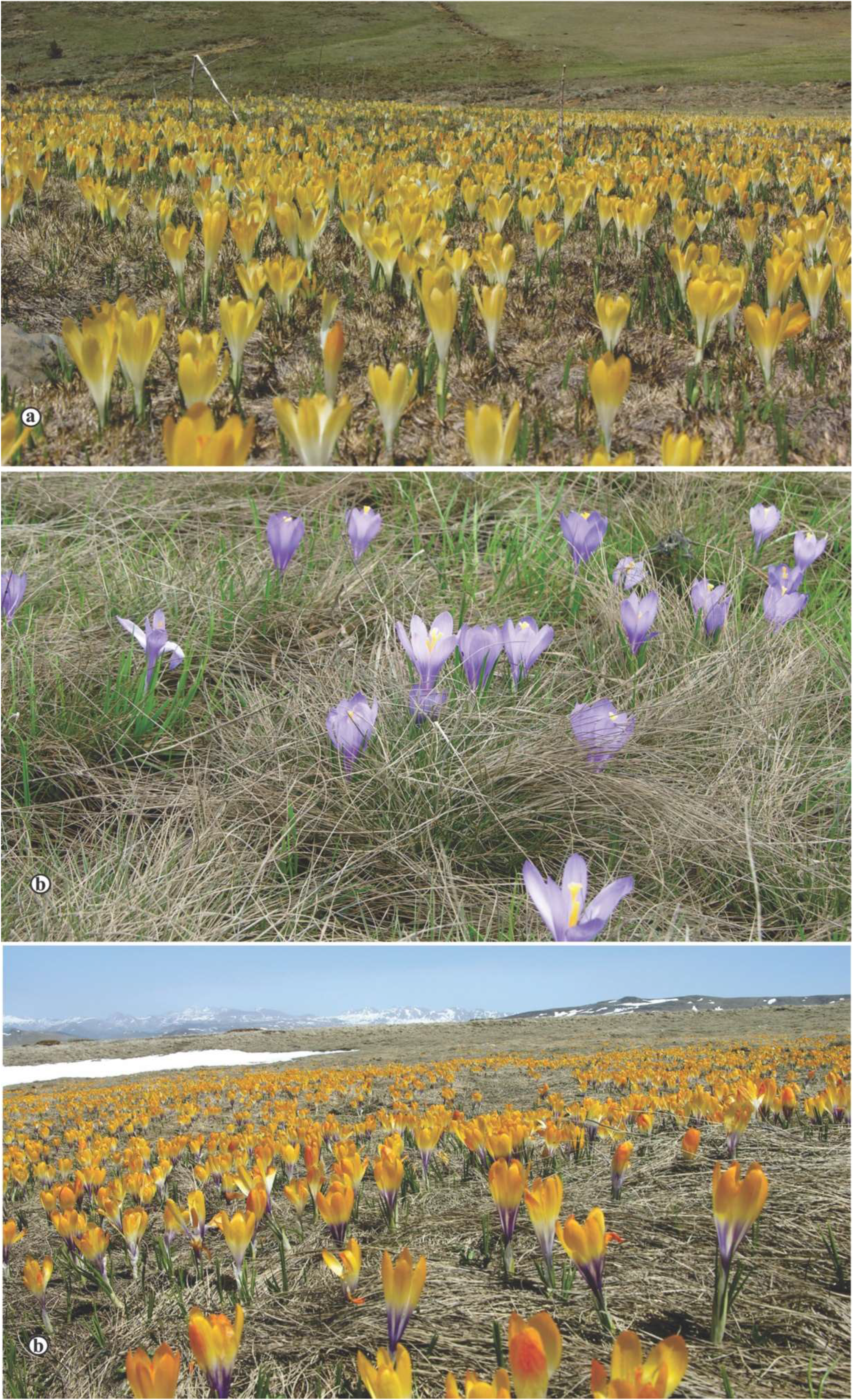
a) *C. albanicus* (Valamara Mt.), b) *C. doerteae* (Liqeni i Madh, Mbas Deja Mt.) and c) *C. scardicus* (Sorokol, Korabi Mt.) in their habitats.

Type:—ALBANIA. Korça district, Maliqi municipality, serpentine alpine pastures and meadows of Valamara Mt, above Strelca village, 06 June 2019, 2070 m, *D. Shuka s*.*n*. (holotype: TIR!; isotypes: GAT0083264-0083267!, ZAGR!).

Bulbous, perennial, 7–12 cm high. Corm flattened-subglobose, 8–14 mm in diameter and 4–7 mm high, covered by several layers of reticulate fibers outside and membranous layers with reticulated fibers inside that come out from under **a thick coriaceous corm-cap** (fig. 9: a, b, c). The reticulate fibers in the upper part of the corm are often irregular, ending in parallel fibers around the stem in a long persistent neck 1.5–2 cm. The outer fibers are thicker (0.1–0.14 mm) than the inner fibers (0.08–0.09 mm).

Cataphylls 3–4, white to greenish above the ground, with light green veins at apex, more distinctly in the upper part of perigone tube. Prophyll present. Bract and bracteole present, subequal to equal, exerted from cataphylls, shorter than the perigone tube, rare equal. Leaves (3–) 4 (–5), present at anthesis, 1.2–2 (–2.2) mm wide and 0.6–1 mm thick, glabrous, shiny green without white median strip above. Leave cross section show a regular “**T**” shape of adaxial surface, and leaf-arms with spongy parenchyma and abaxial part of it. Abaxial part of leave is 0.8–1.2 mm wide or as much as 2/3 of the width of the leaves or adaxial part.

Perigone tube white, 2.5–5 cm. Throat white or yellowish both inside and outside, which becomes white going towards the base of perigone segments, glabrous. Flowers solitary, very rarely 2, yellow to orange towards of the segment’s apex, rarely completely white or yellow. Outer perigone segments slightly longer than inner ones or equal. Segments 3–4 (–4.3) × 0.6– 1.2 cm, oblanceolate or elliptical, obtuse to subacute.

Filaments white 10–15 (–17) mm long, glabrous, dilated at base; anthers yellow, 10–15 mm long included the lobes, equal or shorter than filaments, pollen yellow. Style (20–) 26–34 (– 40) mm long, glabrous, equal or longer than anthers, white in 2/3 of lower part, yellow and orange towards the apex, with usually simple three orange expanded branches, 3–7 mm long, branching position in relation to the anthers mostly in the upper third of the anthers and above of them. Stems in ripen capsules (6–)8 × 12 cm above the ground, capsule 12–16 × 3–5 mm, ellipsoid. Seeds 2.5–4 × 1.5–2 mm, dark reddish-brown at maturity.

### Habitat and Distribution

*C. albanicus* occurs only in the damp, semi-wet or semi-dried pastures in the ophiolitic areas of Valamara Mts, mostly belongs to alkaline transition mires and quaking bogs. The growth habitats of *C. albanicus* lies in alpine part of the serpentine massif, which has a length of about 25 km and represents the northernmost edge of the northern Pindos. The habitat is characterized by deep layer of soil or peat with pH 7.6-8 and very low nutrients (total N and P), but rich with heavy metals such as Cr, Fe, Mg, Ni, Pb (S∤omka et al. 2015). However, these measurements were done in in dried pastures and not in the wet meadows, which is the main habitat of *C. albanicus*. The vegetation of alluvial damps, semi-wet or semi-dried pastures is dominated by associations *Veratrum album-Carex* sp. div., *Nardetum strictae-Festuca* sp. div. (Buzo 1991) and *Carici sempervirentis*-*Seslerietum coerulantis* (Fanelli et al. 2018) with the following dominant species: *Veratrum album, Carex* spp., *Narthecium scardicum, Nardus stricta, Festuca panciciana, Cirsium appendiculatum, Pinguicula hirtiflora* and *P. balcanica, Caltha palustris, Geum coccineum, Hieracium pilosella* etc. Along the water courses or around them, it was companied with *Centaurea pindicola, C. vlachourum, Heliosperma pusillum* subsp. *albanicum Silene parnassica* subsp. *pindicola, Viola albanica* and *V. ducadjinica*, indicators of serpentine substrates (Shuka 2009, Shuka & Tan 2009, Miszczak et al. 2022).

### Other specimen examined of *Crocus albanicus*

—ALBANIA. Gramshi municipality, Lenie village, abave Black Lake, in a semi-wet serpentine meadow in the Valamara Mt range, 2070 m, 29 May 2020, Leg. *D. Shuka s*.*n*. (herb. Shuka), TIR!).

—*loc. ibid*., above Grabova village, in serpentine pastures of the Valamara Mt, 2170 m, 29 May 2020, Leg. *D. Shuka s*.*n*. (herb. Shuka), TIR!).

—Pogradeci municipality, Lukova Lakes, in wet serpentine places around the lower lake, 1730 m, 19 May 2021, Leg. *L. Shuka s*.*n*. (herb. Shuka), TIR!).

### Specimens examined of *C. scardicus*

—ALBANIA. Kukësi municipality, Kallabak Mt, in siliceous grasslands, 1700-1900 m, 08 June 2020, *D. Shuka* & *B. Hallaçi s*.*n*. (herb. Shuka).

— Peshkopia municipality, pastures of Sorokol, North of Korabi Mt, in siliceous grasslands, 1800-2200 m, 09 June 2020, *D. Shuka* & *B. Hallaçi s*.*n*. (herb. Shuka).

—KOSOVA. Dragashi municipality, alpine pastures and meadows above Brod and Restelica villages, at the foot of Vraca Mts, 2100 m, 15 June 2013, *L. Shuka s*.*n*. (herb. Shuka).

—*loc. ibid*., in siliceous grasslands on the southwestern part of Restelica village, 1900 m, 07 June 2020, *D. Shuka* & *B. Hallaçi s*.*n*. (herb. Shuka).

## Remarks and conclusions

All four series *Scardicii* species including the two newly described ones, can be distinguish by their morphological characters, are genetically differentiated from each other, are growing at different habitat types and/or mountains, which differ in geomorphology. Aside from geographical isolation, adaption might have been a driving factor in species differentiation within the series *Scardici*.

Series *Scardici* crocuses are narrow endemics within a restricted distribution area in the mountains of the Western Balkans. Such alpine species are highly threatened by climate change causing a decrease of snow cover and precipitation and therefore, increasing drought conditions (Knez et al. 2022). Additionally, the ongoing development and extension of tourism in the mountainous regions are threating the habitats. Therefore, population dynamics and species should be monitored and assessed.

## Acknowledgements

Fieldwork in Valamara Mt. for D. Shuka was supported by a grant from AKKSHI (National Agency for Scientific Research and Innovation). I (D.S) am very indebted to Mr. L. Firaj for accompanying me in the field trips carried out in Valamara Mt.

We like to thank I. Faustmann, C. Koch, B. Kraenzlin and P. Oswald for help with lab work. This work was supported by funding from Deutsche Forschungsgemeinschaft (DFG) through grant HA7550/2 to D.H.

## Notes

### Competing Interest Statement

The authors have declared no competing interest.

