## supplementary table for "Two new species of *Crocus* series *Scardici* from the Western Balkans"

| **Gene family** | **Region name** | **Primer name** | **Primer sequence (5’-3’)** | **Amplicon size in bp** | **PCR annealing temperature**  **(Phusion Taq)** |
| --- | --- | --- | --- | --- | --- |
| *ORTHO04x5M007634* | embo | embo_f | CTGATAGACATTGGATATTTCCC | ~700 | 60 °C |
|  |  | embo_r | AAGTTTCTCAGGGAACATCGG |  |  |

**Table**  PCR and primer information of the newly applied marker.
